# The Listening Zone of Human Electrocorticographic Field Potential Recordings

**DOI:** 10.1101/2021.10.22.465519

**Authors:** Meredith J McCarty, Oscar Woolnough, John C. Mosher, John Seymour, Nitin Tandon

## Abstract

Intracranial electroencephalographic (icEEG) recordings provide invaluable insights into neural dynamics in humans due to their unmatched spatiotemporal resolution. Yet, such recordings reflect the combined activity of multiple underlying generators, confounding the ability to resolve spatially distinct neural sources. To empirically quantify the listening zone of icEEG recordings, we computed the correlations between signals as a function of distance (expressed as full width at half maximum; FWHM) between 8,752 recording sites in 71 patients implanted with either subdural electrodes (SDE), stereo-encephalography electrodes (sEEG), or high-density sEEG electrodes. As expected, for both SDE and sEEG electrodes, higher frequency signals exhibited a sharper fall off relative to lower frequency signals. For broadband high gamma (BHG) activity, the mean FWHM of SDEs (6.6 ± 2.5 mm) and sEEGs in gray matter (7.14 ± 1.7 mm) was not significantly different, however the FWHM for low frequencies recorded by sEEGs was 2.45 mm smaller than SDEs. White matter sEEG electrodes showed much lower power for frequencies 17 to 200 Hz (q < 0.01) and a much broader decay (11.3 ± 3.2 mm) than gray matter electrodes (7.14 ± 1.7 mm). The use of a bipolar referencing scheme significantly lowered FWHM for sEEG electrodes, as compared with a white matter reference or a common average reference. These results outline the influence of array design, spectral bands, and referencing schema on local field potential recordings and source localization in icEEG recordings in humans. The metrics we derive have immediate relevance to the analysis and interpretation of both cognitive and epileptic data.

## Introduction

Invasive neural recordings provide a unique window into human cognition. Over the last several decades, intracranial field potential recordings have yielded profound insights into a variety of neural systems including speech production (Cogan et al. 2014; Pasley et al. 2012), auditory processing (Miller et al. 2021), language (Conner et al. 2019; Forseth et al. 2018), visual perception (Martin et al. 2019), motor control (Salari et al. 2019), decision making (Bartoli et al. 2018), emotion (Guillory and Bujarski 2014), and memory (Derner et al. 2018; Foster et al. 2012). An array of electrode designs and recording scales are now being implemented and ongoing progress in neuroengineering is yielding rapid advances in electrode design. The gap between what recording scale is technologically possible and that which is optimal for understanding the neurobiology of cognition, epilepsy or to provide inputs for brain machine interfaces, remains unknown (Marblestone et al. 2013; Pesaran et al. 2018). Answers to these questions, especially the optimal form factor required to resolve spatially distinct sources within the complex electric field landscape of the brain will influence the design of newer recording interfaces (Cybulski et al. 2015).

The uncertainty of reconstructing the spatial and temporal sources based on multi-electrode field potentials - the inverse source problem (Herreras 2016; Pesaran et al. 2018) is a direct consequence of the imperfect resolution of recording electrodes and the source properties of the electric field landscape. While the complex geometry of single neurons makes the precise modeling of even one neuron’s activity in isolation difficult to model (Nunez and Srinivasan 2005), the field potential at any recording electrode is an aggregate of quasi-synchronously active dipoles from a multitude of spatially distributed neural sources (Buzsáki et al. 2012; Łęski et al. 2013). Not all neurons contribute to this electric field landscape at any given instant, and different patterns of neural activity may generate similar field potential measures depending on the distance and the density of recording sites. The neural tissue that comprises this electric field landscape is itself heterogenous, with conductivity and dielectric constants that vary based on cell packing density and cortical location (Bingham et al. 2020; Howell and McIntyre 2016; Nunez and Srinivasan 2005).

At the resolution currently provided by macroelectrodes used for human intracranial electroencephalographic (icEEG) recordings, the measured field potential activity is not a direct measure of the activity of local cell assemblies, but rather a larger-scale measure of activity conducted through neural space. This volume conduction can lead to linear relationships between simultaneously recorded signals at neighboring electrodes, and it is hard to disentangle whether high levels of correlated activity between two electrodes are due to underlying neural dynamics (such as common input to both regions) or due to volume conduction of voltage from neighboring regions (Kellis et al. 2016). To resolve this, we define and quantify volume conduction as the instantaneous signal correlation at zero-time lag between electrode pairs, which quantifies common activity due to volume conduction. The lower the instantaneous correlation between electrodes, the lower the signal redundancy of each electrode’s listening zone and the greater its uniqueness. Determining the optimal spacing and location of electrodes to not only minimize signal redundancy, but to also capture separable field potential recordings is a pivotal hurdle for understanding and optimizing invasive field potential recordings in humans (Cybulski et al. 2015).

To investigate the ability of multiple clinically used electrode types in resolving spatially distinct activity, we compared task-related cross-correlations in activity across subdural electrodes (SDE), stereo-electroencephalography electrodes (sEEG), and high-density sEEG (hdsEEG) electrodes in patients undergoing monitoring for the localization of medically intractable epilepsy. We analyzed the impact that referencing strategy, electrode location, and frequency components of the signal have on signal redundancy and the influence this could have on neural array design.

## Materials and Methods

### Participants

71 patients (33 female, 18-65 years) participated in this research after providing written informed consent. All participants were semi-chronically implanted with intracranial electrodes for the localization of pharmaco-resistant epilepsy. All experimental procedures were reviewed and approved by the Committee for the Protection of Human Subjects (CPHS) of the University of Texas Health Science Center at Houston as Protocol Number HSC-MS-06-0385.

### Electrode Implantation and Data Recording

Data were acquired from either subdural grid electrodes (SDEs; 18 patients), stereotactically placed depth electrodes (sEEGs; 47 patients) or high-density depth electrodes (hdsEEGs; 6 patients) (Figure 1C,D). SDEs were subdural platinum-iridium electrodes embedded in a silicone elastomer sheet (PMT Corporation; top-hat design; 3mm diameter cortical contact), surgically implanted via a craniotomy (Conner et al. 2011; Pieters et al. 2013; Tandon 2012; Tong et al. 2020). sEEGs were implanted using a Robotic Surgical Assistant (ROSA; Medtech, Montpellier, France) (Rollo et al. 2020; Tandon et al. 2019). Each sEEG probe (PMT corporation, Chanhassen, Minnesota) was 0.8 mm in diameter and had 8-16 electrode contacts. For the standard sEEG electrodes, each contact was a platinum-iridium cylinder, 2.0 mm in length and separated from the adjacent contact by 1.5 - 2.43 mm. Each patient had 12 - 20 sEEG probes implanted. For hdsEEG electrodes, contacts were 0.5 mm in length and separated from the adjacent contact by 0.5 mm. Each patient had 1 - 4 hdsEEG probes implanted. Following surgical implantation, electrodes were localized by co-registration of pre-operative anatomical 3T MRI and post-operative CT scans in AFNI (Cox 1996). Electrode positions were projected onto a cortical surface model generated in FreeSurfer (Dale et al. 1999), and displayed on the cortical surface model for visualization (Pieters et al. 2013). Intracranial data were collected during research experiments starting on the first day after electrode implantation for sEEGs and two days after implantation for SDEs. Data were digitized at 2 kHz using the NeuroPort recording system (Blackrock Microsystems, Salt Lake City, Utah), imported into Matlab, initially referenced to the white matter electrode used as a reference for the clinical acquisition system and visually inspected for line noise, artifacts and epileptic activity. Electrodes with excessive line noise or localized to sites of seizure onset were excluded. Trials contaminated by inter-ictal epileptic spikes were discarded.

**Figure 1.**
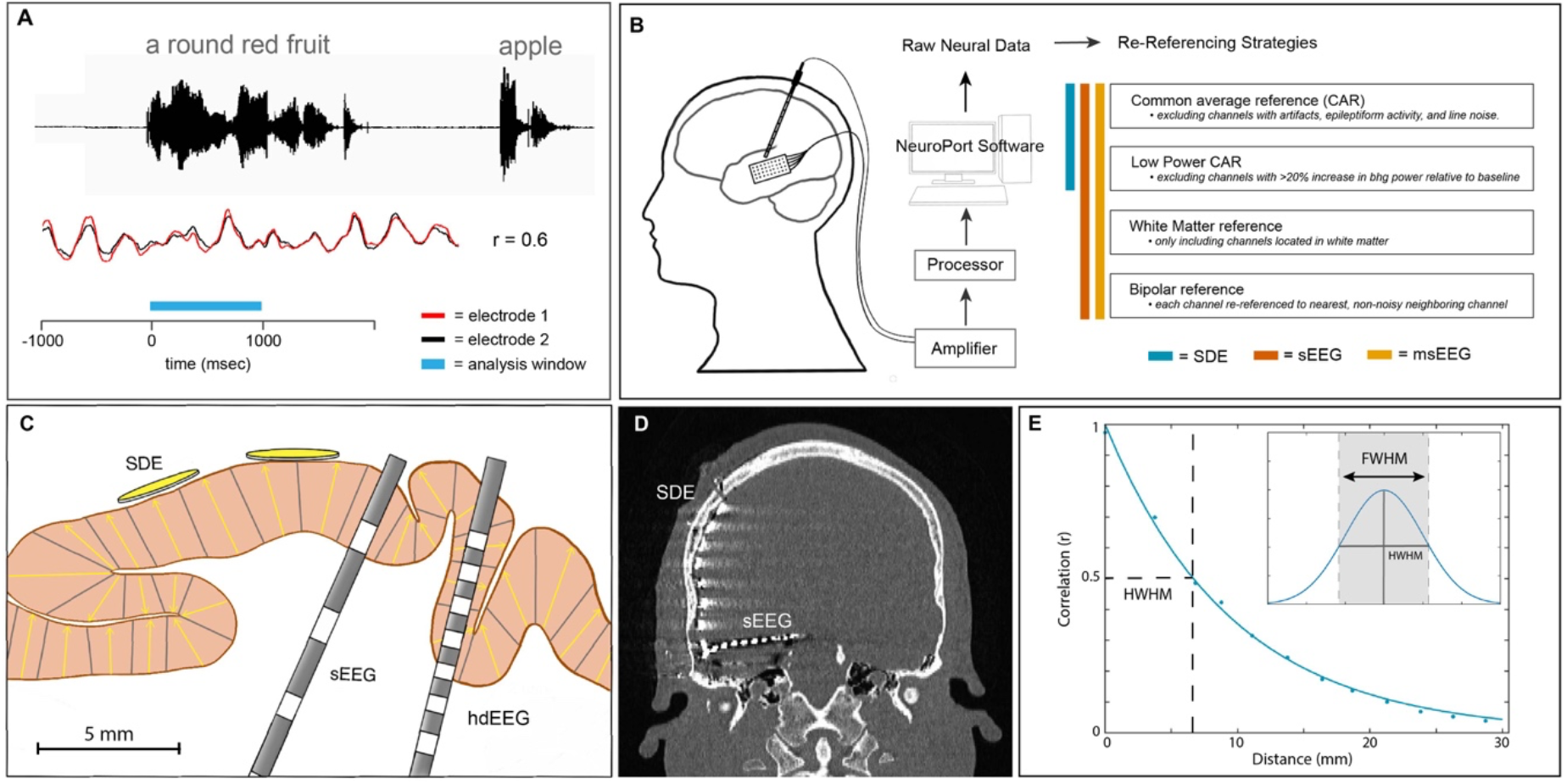
Experimental Design. (**A**) Schematic representation of the auditory naming to definition task. Colored bar indicates task-related analysis window (blue; 0 to 1000ms) during which cross-correlation (r) is calculated between the waveforms of two exemplar neighboring electrodes (red and black; exemplar traces). (**B**) Schematic representation of the neural data acquisition and re-referencing strategies. (**C**) Schematic representation of the three electrode scales analyzed: subdural electrodes (SDE) 3 mm diameter disc, stereo-electroencephalography (sEEG) electrodes 2-mm long ring, and high-density sEEG (hdsEEG) electrodes 0.5-mm long ring. sEEG and hdsEEG contacts are depicted in grey. Yellow arrows depict dipole orientation within pictured cortical gray matter. (**D**) Representative computed tomography (CT) scan of a patient with concurrently implanted SDE and sEEG electrodes. (**E**) Example of full width at half maximum (FWHM) calculation. The correlation coefficient was measured using the raw voltage of every combination of electrode pairs within 30mm of each other, for each frequency range. Correlation values were fit with an exponential decay function. Half width at half maximum (HWHM) correlation was measured from this exponential decay function and doubled to generate the FWHM value for each condition.

### Signal Analysis

Across all 75 patients, a total of 2,546 SDE, 8,493 sEEG, and 204 hdsEEG electrode contacts were implanted. Of these, 704 SDE, 1,736 sEEG, and 51 hdsEEG were excluded due to proximity to the seizure onset zone, frequent inter-ictal epileptiform spikes or line noise. The remaining electrodes included were: 1,842 SDE, 6,757 sEEG, and 153 hdsEEG electrodes. Analyses were performed by bandpass filtering raw icEEG data from each electrode into 5 frequency bands (Theta, 4-8 Hz; Alpha, 8-15 Hz; Beta, 15-30 Hz; Narrowband Gamma, 30-60 Hz; Broadband High Gamma, 70-150 Hz). Following the removal of line noise (zero-phase 2^nd^ order Butterworth band-stop filters), band-limited voltage traces were obtained (zero-phase 3^rd^ order Butterworth bandpass filters).

### Referencing and Re-referencing strategy

During the recording session, a non-noisy clinical hardware reference electrode located in white matter was used as the reference electrode. For analysis, recordings were re-referenced using one of the following schemes (Figure 1B):

*Common average reference (CAR):* Offline, raw data were visually inspected and electrodes exhibiting electrical noise or epileptiform artifacts were excluded from the common average. Neural data was then re-referenced to the average of all remaining electrodes that were included in this CAR.
*Low-Power CAR:* Broadband high gamma activity (70 – 150 Hz) was extracted for each time series (using the original clinical reference) using a frequency domain Hilbert transform and the percentage change in power was measured relative to a baseline time window of −500 to −100 ms before stimulus onset. If the percentage change in mean power was less than 20%, electrodes were included in the low-power CAR signal averaging.
*White Matter referencing:* We identified all sEEG and hdsEEG electrodes located in white matter, gray matter, and cerebrospinal fluid (CSF) based on their position relative to their FreeSurfer surfaces and included all white matter located electrodes.
*Bipolar referencing:* For the bipolar re-referencing, each electrode on the sEEG and hdsEEG probes was re-referenced to its closest neighboring non-noisy electrode located on the same probe. Electrodes on the end of the probe or whose nearest neighboring electrode was noisy were excluded from the analysis.

### Experimental Design and Statistical Analyses

#### Experimental Task

All patients participated in an auditory naming-to-definition task (Figure 1A) (Forseth et al. 2018), producing single word responses to an auditory presented definition. 70+ auditory stimuli (mean 87) were presented to each patient using stereo speakers (44.1 kHz, 15” MacBook Pro 2015) (Forseth et al. 2018). Stimuli had an average duration of 1970 ± 360 ms, and an inter-stimulus interval of 5000 ms.

The time period of interest for this analysis was from 0 to 1000 ms following auditory stimulus onset.

#### Full width at half-maximum (FWHM) measure

To compare correlation between electrode pairs over distance, we calculated the full width at half maximum (FWHM) correlation. We first identified all non-noisy pairs of electrodes that were less than 30mm from each other (in Euclidean distance). Pairwise Pearson’s correlation was calculated between the band-limited voltage traces for all electrode pairs for each trial (Figure 1A). This correlation value was then averaged across all trials to return one correlation value for each electrode pair and frequency range. A decay function was fit to the absolute values of the correlations within each individual patient (Figure 1E). The decay function was defined as *r* = (1 – *β*)*^d^*, where the correlation *r* decayed based on the decay factor *β* and the distance *d*. The decay factor was optimized using a least-squares fit. From this decay function, we extracted the distance at which the correlation equaled 0.5 - half the theoretical maximum correlation (half width at half maximum; HWHM). The HWHM value was doubled to generate the FWHM value for each condition (Figure 1E). For visualization purposes, the absolute values of these correlations for each patient were binned based on Euclidean distance into 2.5mm bins.

#### Validation on Simulated Data

Simulated timeseries data were created using the neural digital signal processing toolbox (Cole et al. 2019). 100 unique power law timeseries were generated in each of the 5 previously described frequency bands of interest with a power-law exponent of −2, a sampling frequency of 2,000 Hz, and a simulation time of 1.5 seconds to account for the removal of filtering edge effects. The Pearson’s correlation coefficient was calculated between each pair of simulated signals to generate the actual correlation measurement. To calculate the reconstructed correlation dataset, timeseries from each frequency range were first combined to generate a summed electric field signal. Analyses were performed by bandpass filtering combined simulated data into the 5 previous described frequency bands using identical methods to the main analysis. Signals were randomly re-paired to create 75 simulated trials, approximately matching experimental conditions (Supplemental Figure 1).

#### Power Spectral Density (PSD) Analysis

Thomson’s Slepian multitaper power spectral density (PSD) estimate of the signal was calculated. Significant differences between power in gray and white matter was calculated with Wilcoxon sign rank tests, corrected for multiple comparisons using a Benjamini-Hochberg false detection rate (FDR) threshold of q<0.01.

#### Linear Mixed Effects (LME) Modelling

Linear mixed effects models were used to incorporate random and fixed-effects into a linear model. Fixed effects in our model were electrode type and frequency band. The random effect in our model was the participant. Electrode type was SDE, sEEG or hdsEEG. Data were assumed to be normal in distribution for statistical comparison.

#### Data Visualization using Raincloud plots

Raincloud plots, incorporating raw data points, probability density, and median, mean, confidence intervals, were utilized to visualize data (Allen et al. 2019). Reported values for each category are median ± interquartile range.

## Results

We utilized a correlation-based analysis to compute the falloff of cross-correlation as a function of distance, between pairs of all non-noisy electrodes regardless of cortical location. We constrained our analysis to task-related neural data, based on prior evidence that the spatial spread of correlated activity is lower during activity as opposed to rest (Muller et al. 2016). Importantly, our analyses compare differences in FWHM across referencing conditions, thereby preserving inter-electrode distance as a variable. By preserving inter-electrode distance in our FWHM measures, we effectively compute a local reduction in correlation, rather than a global reduction, as is captured in other distance-averaged correlation comparisons.

### Effect of electrode scale and signal frequency on listening zone

We first compared the decay function (indexed by the FWHM) for SDE vs sEEG electrodes in gray matter, to determine whether a subdural or intracortical location of the icEEG electrode significantly influences the listening zone (Figure 2). To compare differences in FWHM across frequency and electrode scale, we used a linear mixed effects (LME) model with fixed effects modeling frequency bands and electrode scale (SDE or sEEG). This model explained a large proportion of the variance of FWHM measures (r^2^ = 0.65). The electrode type had a significant effect on FWHM (t(321) = −4.5, β = −2.4, P < 0.001, 95% CI −3.5 to −1.4), which was 2.45 mm smaller for sEEG electrodes than for SDE electrode pairs, when comparing across all frequency ranges. The FWHM of the decay was smaller as frequency increased (LME: t(321) = −16.0 β = −1.8, P < 0.001, 95% CI −2.0 to −1.6) and there was a significant interaction between frequency and electrode type (t(321) = 3.2, β = −0.42, P = 0.001, 95% CI 0.17 to 0.68) indicating that the spatial extent of correlation is significantly dependent on frequency and electrode scale.

**Figure 2.**
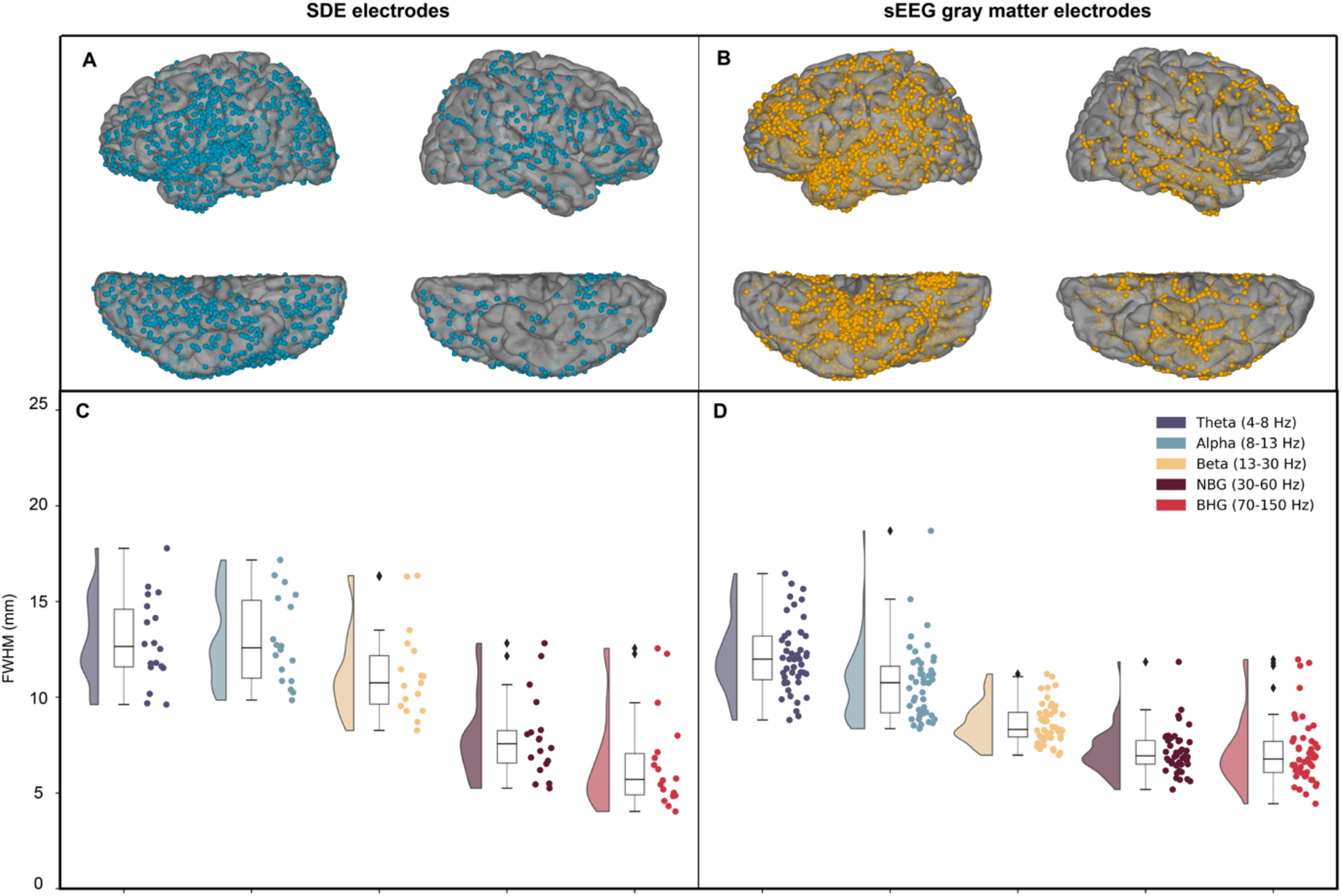
Across-electrode differences in correlation over distance. Coverage map of locations of SDE electrodes (**A**; 18 patients, 1,842 electrodes, 37,272 electrode pairs) and gray matter located sEEG electrodes (**B**; 47 patients, 2,916 electrodes, 47,522 electrode pairs). Average full width at half maximum (FWHM) was calculated and plotted for each patient for SDE (**C**) and sEEG (**D**) electrode pairs. Abbreviations: narrowband gamma (NBG), broadband high gamma (BHG).

For BHG alone, electrode type did not have a significant effect on FWHM (t(34) = 1.42, β = 1.07, P = 0.17, 95% CI −0.5 to 2.6). The mean FWHM in BHG for SDE electrodes (6.6 ± 2.5 mm) was slightly lower than for gray matter located sEEG electrodes (7.14 ± 1.7 mm), however this difference was not significant.

### Location dependence of sEEG electrode listening zone

SDEs sit on the cortical surface, proximal to local field generators, whereas many individual sEEG electrodes are located within white matter, distant from the cortical surface and measuring far field potentials. Thus, the physical location of sEEG electrodes could present potentially confounding correlation measures across distance. An LME model with fixed effects modeling frequency and electrode location (white matter or gray matter located sEEGs) explained a large proportion of the variance in FWHM measures (r^2^ = 0.78). sEEG electrodes located in gray matter had a much smaller FWHM (8.3 mm lower) compared to those located in white matter (LME: t(466) = −17.3, β = −8.3, P < 0.001, 95% CI −9.2 to −7.3) (Figure 3, Supplementary Figure 2). Additionally, the interaction between FWHM and frequency range significantly depended on electrode location (t(466) = 6.6, β = 0.95, P < 0.001, 95% CI 0.67 to 1.2) with low frequencies showing a broader listening zone in white matter electrodes. For theta frequencies, the mean FWHM for sEEG electrodes located in white matter was 20.2 ± 4.3 mm, whereas the mean FWHM for gray matter sEEG electrodes was 12.1 ± 1.8 mm.

**Figure 3.**
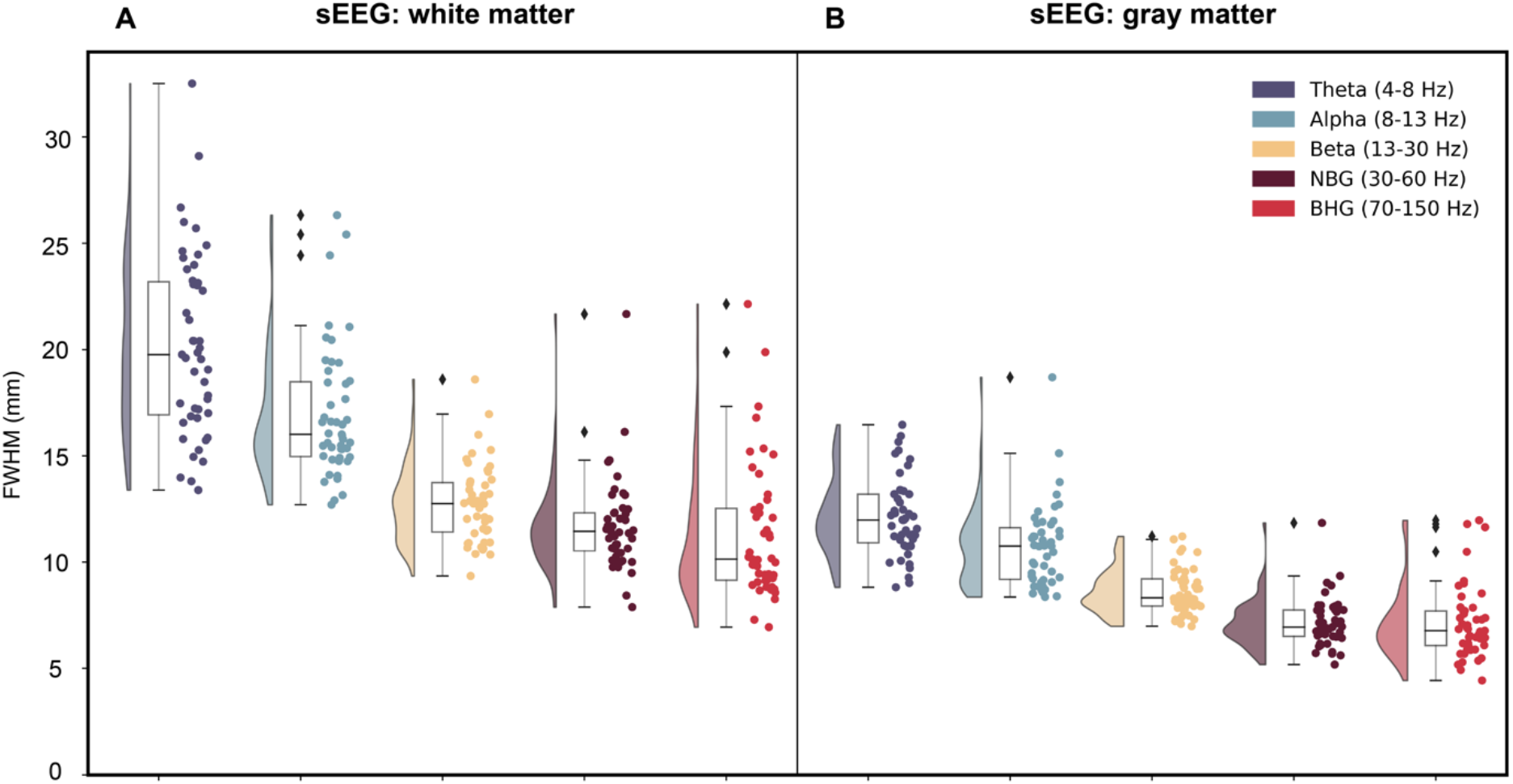
Anatomical location of sEEG contacts in gray or white matter significantly influences full width at half maximum (FWHM) correlation measures. Raincloud plots depicting FWHM values for each patient in each frequency range for all pairs of white matter located (**A**; 2,649 electrodes; 43,957 electrode pairs) and gray matter located (**B**; 2,916 electrodes; 47,522 electrode pairs) pairs. Abbreviations: narrowband gamma (NBG), broadband high gamma (BHG).

When comparing the effect of electrode location on BHG activity, an LME model with fixed effects modeling electrode location explained a large proportion of the variance of the FWHM measures (r^2^ = 0.85). For BHG frequencies, the mean FWHM for sEEG electrodes located in white matter was 11.3 ± 3.2 mm, whereas the mean FWHM for sEEG electrodes located in gray matter was 7.14 ± 1.7 mm. For the BHG band, electrode location did have a significant effect on FWHM of signal correlation decay (t(92) = −13.5, β = −4.2, P < 0.001, 95% CI −4.8 to −3.6). Of course, there is not much power in white matter recordings and these correlations may be higher given these lower amplitude signals. To assess this, we compared mean power spectral density (PSD) plots for sEEG electrodes located in white matter or gray matter, demonstrating the much lower power in white matter sEEG electrodes (for all frequencies 18 to 200 Hz; q < 0.01) (Supplementary Figure 3).

### Referencing strategies for SDE and sEEG electrodes

Next, we examined the influence of referencing schemes on measured correlation. Based on evidence that referencing strategies can eliminate or increase spurious correlation between recording electrodes (Li et al. 2018), we compared several commonly used referencing schemes; common average reference (CAR), low-power CAR, white matter referencing, and bipolar referencing, across SDE and gray matter located sEEG electrode pairs (Figure 4).

**Figure 4.**
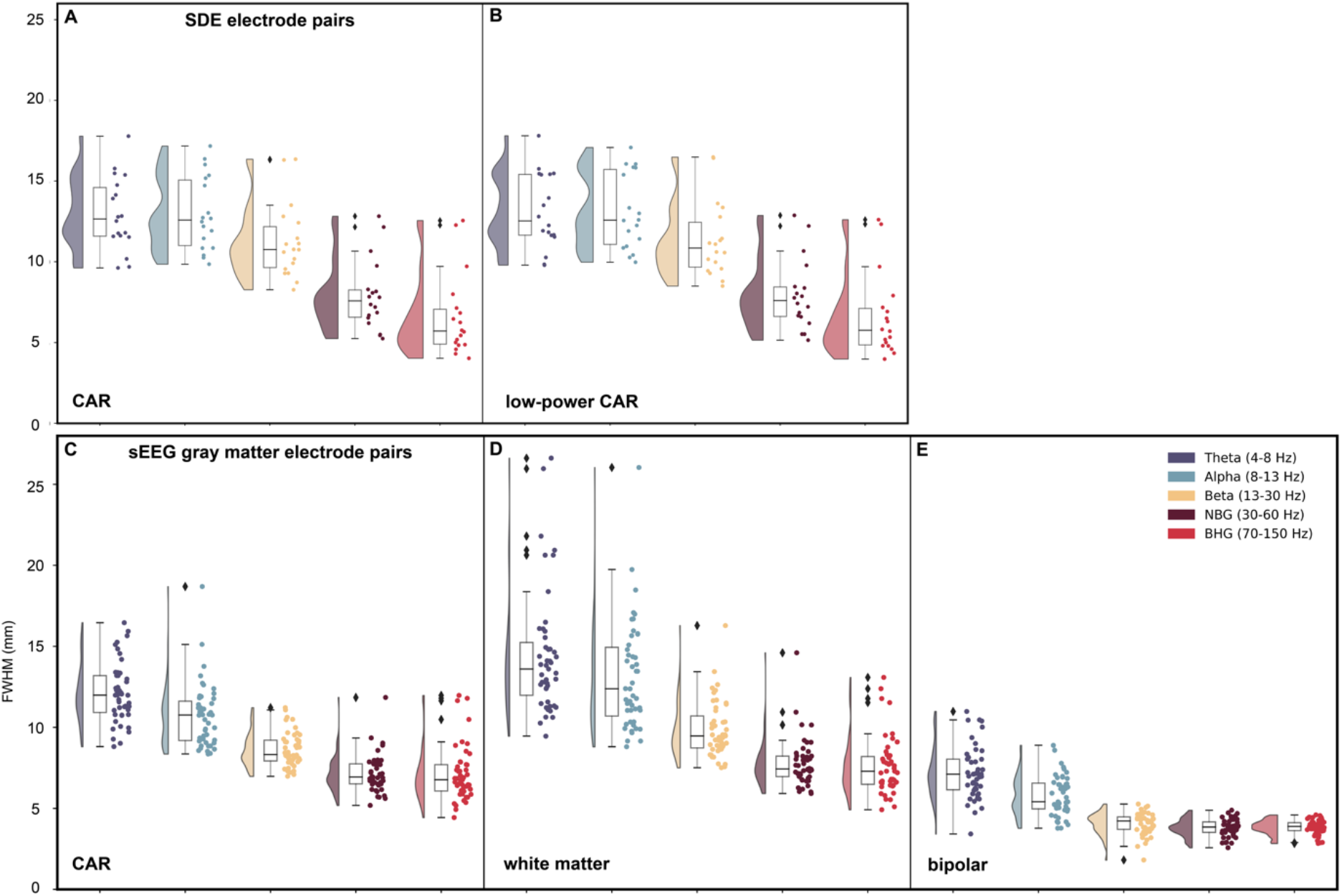
Referencing scheme comparison across SDEs and gray matter located sEEGs. Average full width at half maximum (FWHM) was calculated and plotted for each patient for SDE (**A-B**; n = 18 patients) and sEEG (**C-E**; n = 47 patients) electrode pairs in each frequency range of interest. For SDE electrode pairs (1,842 electrodes; 37,272 electrode pairs), average FWHM was compared using either common average reference (CAR) (**A**) or low-power CAR scheme (**B**). For gray matter located sEEG electrode pairs (2,916 electrodes; 47,522 electrode pairs), average FWHM was compared using either CAR (**C**), white matter (**D**), or bipolar referencing schemes (**E**). Abbreviations: narrowband gamma (NBG), broadband high gamma (BHG).

A two-way ANOVA was conducted comparing effects of referencing scheme and frequency range on FWHM measures. For SDE electrode pairs, there was no significant interaction between the referencing scheme and the frequency band on FWHM measures (F(4,170) = 0.01, P = 0.99). There was a significant effect of frequency (F(4,170) = 57.96, P < 0.001) on FWHM, but no significant effect of referencing scheme (F(1,170) = 0.07, P = 0.79). For BHG activity, SDEs showed a correlation decay of 6.6 ± 2.5 mm FWHM for the CAR scheme (Figure 4A), and 6.6 ± 2.6 mm FWHM for the low-power CAR scheme (Figure 4B). For BHG frequency specifically, a two-way ANOVA showed no significant effect of referencing scheme on FWHM for SDE electrodes (F(1,35) = 5.5 x 10^-5^, P = 0.99). For theta activity, SDEs showed a correlation decay of 13.0 ± 2.3 mm FWHM for the CAR scheme (Figure 4A), and 13.1 ± 2.3 mm FWHM for the low-power CAR scheme (Figure 4B).

For gray matter located sEEG electrode pairs, a two-way ANOVA showed a significant effect of referencing type (F(2,690) = 586.48, P < 0.001) and frequency (F(4,690) = 207.83, P < 0.001) on FWHM values. There was a significant interaction between frequency and referencing scheme on FWHM values (F(8,690) = 9.92, P < 0.001). For BHG activity, sEEGs showed a correlation decay of 7.14 ± 1.7 mm FWHM for the CAR scheme (Figure 4C), 7.62 ± 1.8 mm for the white matter referencing scheme (Figure 4D), and 3.83 ± 0.45 mm for the bipolar referencing scheme (Figure 4E). For BHG frequency specifically, a two-way ANOVA showed a significant effect of referencing scheme on FWHM for sEEG electrodes (F(2,140) = 94.4, P < 0.001). For theta activity, sEEGs showed a correlation decay of 12.1 ± 1.8 mm FWHM for the CAR scheme (Figure 4C), 14.4 ± 3.8 mm for the white matter referencing scheme (Figure 4D), and 7.19 ± 1.6 mm for the bipolar referencing scheme (Figure 4E).

### Listening zone of high-density sEEG (hdsEEG) electrodes

The final group analysis compared pairwise correlation between hdsEEG electrodes (6 patients; 153 electrodes) across referencing scheme. These electrodes were cylinders of 0.5 mm length as compared to 2 mm contacts in standard sEEGs. (Figure 5A). For broadband gamma activity, hdsEEG electrode pairs (CAR) had a mean FWHM of 6.5 ± 1.4 mm relative to the FWHM for gray matter located sEEG electrode pairs (7.14 ± 1.7 mm) and the SDE electrode pairs (6.6 ± 2.5 mm). For BHG frequency specifically, a two-way ANOVA showed no significant effect of referencing scheme on FWHM for hdsEEG electrodes (F(2,17) = 0, P = 0.998). For theta activity, hdsEEG electrode pairs (CAR) had a mean FWHM of 17.3 ± 6.7 mm relative to FWHM for gray matter sEEG electrode pairs (12.1 ± 1.8 mm) and SDE electrode pairs (13.0 ± 2.3 mm).

**Figure 5.**
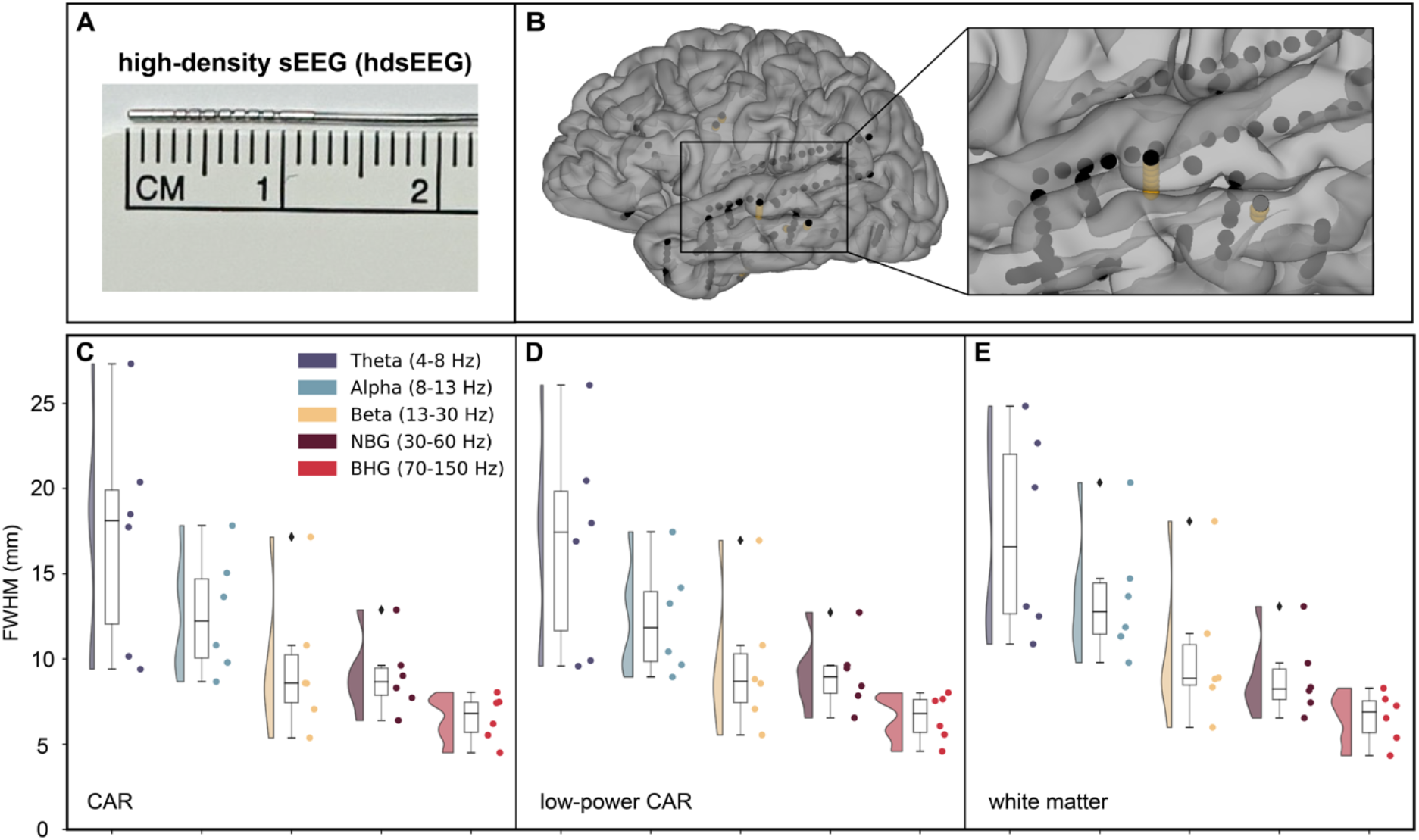
Referencing scheme comparison across hdsEEG electrodes. hdsEEG electrodes have 0.5 mm in length electrodes (**A**) and an exemplar hdsEEG electrode implanted in one patient is shown in yellow (**B**). Average full width at half maximum (FWHM) was calculated and plotted for each patient for hdsEEG electrode pairs in each frequency range of interest. For hdsEEG electrode pairs (6 patients; 153 electrodes; 1,967 electrode pairs), average FWHM was compared using either CAR (**C**), low-power CAR (**D**) or white matter referencing schemes (**E**). Abbreviations: narrowband gamma (NBG), broadband high gamma (BHG).

We compared CAR (Figure 5C), low-power CAR (Figure 5D), and white matter (Figure 5E) referencing schemes for hdsEEG electrode pairs. A two-way ANOVA was conducted comparing effects of referencing scheme and frequency range on FWHM measures. There was no significant interaction between the effects of referencing scheme and frequency range on FWHM measures (F(8,75) = 0.04, P = 1.0). Higher frequencies had significantly lower FWHM than lower frequencies (F(4,75) = 19.59, P < 0.001), and referencing scheme had no effect on FWHM measures (F(2,75) = 0.11, P = 0.90). The location of hdsEEG electrodes in white or gray matter had no significant effect on correlation over distance measures (Supplementary Figure 4).

### Methodological validation on simulated neural data

To validate our analysis pipeline, we also analyzed simulated neural time series data (Cole et al. 2019) in the known correlation values between narrowband signals (Supplementary Figure 1). We found no significant differences between the actual correlations and those reconstructed after running the data through our analysis pipeline.

## Discussion

We have systematically quantified the influence of electrode type, reference scheme and frequency band on the ability to dissociate sources at different scales in human icEEG recordings. Our work shows significant differences in the listening zone across electrode type and frequency band, with SDEs exhibiting the largest listening zone on average relative to sEEGs or hdsEEGs for low frequencies. When considering only high gamma activity, the listening zone was comparable across SDE and sEEG electrodes. The location of sEEG electrodes significantly influenced FWHM measures, with sEEG electrodes located in white matter exhibiting lower power and greater FWHM values than those located in gray matter. There is a significant interaction between spectral band and FWHM for all electrode types, with high frequency gamma signals exhibiting faster fall off of correlation over distance relative to lower frequency signals. Referencing schema only had a significant effect on FWHM measures for sEEG electrodes, with bipolar referencing generating significantly lower FWHM measures as compared with common average or white matter referencing.

### Influence of electrode type on FWHM measures

The location of each electrode, whether atop the cortical surface (SDEs) or intracortically located (sEEGs) led to substantive differences in the listening zone of these electrodes. Across all frequencies, SDEs had broader spread of correlation over distance, with an average FWHM 2.45 mm greater than sEEGs, indicating a more local listening zone for sEEG electrodes. Importantly, the mean FWHM for broadband high gamma alone was not significantly different between SDE (6.6 ± 2.5 mm) and sEEGs (7.14 ± 1.7 mm), indicating a preserved locality of BHG across electrode scale.

The hdsEEG electrodes explored in this analysis are parts of hybrid probes along with traditional sEEG electrodes, and the junction between the conducting and non-conducting edges of the electrode are negligible, as the diameter of the probe is identical across type. While SDE, sEEG, and hdsEEG electrodes have roughly similar impedance, their spacing and location relative to the cortical sources varies significantly. hdsEEG electrode pairs had mean FWHM of 6.5 ± 1.4 mm for BHG, exhibiting the most local listening zone for correlation over distance, albeit in a smaller patient cohort with less electrode pairs in each patient than the SDE and sEEG comparisons.

While there is no consensus on the effect that various recording electrodes have on potential distribution, an electrode’s surface impedance, distance from the source, and source strength all affect source localization (Ellenrieder et al. 2021; Næss et al. 2021; Vermaas et al. 2020). In addition to the inverse problem, a physical factor confounding recorded signals is that electrodes act as capacitors, and their size and impedance (the degree of resistance and reactance with surrounding electric potentials) impacts the resolution of the data (Hnazaee et al. 2020; Moffitt and McIntyre 2005). Fitting with the literature, our analyses reveal differences in the listening zone across electrode types. The SDE, sEEG, and hdSEEG electrodes examined here all have varying electrode size, orientation, spacing, and cortical location, which introduces distinct physical differences in resolution, especially when considering activity in lower frequency ranges.

### Interaction between spectral band and FWHM measures

When examining properties of volume conduction, we found a significant interaction with the spectral band of the filtered EEG signal. SDE electrode pairs exhibited a significant falloff in FWHM as frequency band of interest increased, with mean FWHM being 13.0 ± 2.3 mm for the theta range, whereas mean FWHM for BHG was 6.6 ± 2.5 mm (Figure 2C). This increased FWHM for SDEs at lower frequencies is consistent with a larger spatial reach of lower frequency potential produced by more extensive neuronal generators, which likely induces common activity across a larger region of neural space than smaller generators of higher-frequency activity (Ellenrieder et al. 2021). This difference in FWHM across frequency range was less robust in gray matter located sEEG electrode pairs, with FWHM being 12.1 ± 1.8 mm for theta and 7.14 ± 1.7 mm for BHG range (Figure 2D). When comparing electrode location, white matter located sEEG electrodes did exhibit more frequency-varying falloff of spatial source over distance relative to gray matter located sEEG electrodes (Figure 3). Across electrode scale, high frequency gamma signals exhibited a faster fall off of correlation values across distance, consistent with a smaller spatial reach of a local, weaker, and less synchronous high frequency gamma signal (Dubey and Ray 2020; Ellenrieder et al. 2021; Łęski et al. 2013). This is concordant with synchronous low frequency activity engaging a larger neural substrate than more focal and transient high-frequency activity (Lachaux et al. 2012; Parvizi and Kastner 2018; Rouse et al. 2016; Torres et al. 2019). Interestingly, this fall off of correlation values at lower frequencies varied across electrode type. The mean FWHM for BHG for gray matter sEEG electrodes (7.14 ± 1.7 mm), SDE electrodes (6.6 ± 2.5 mm), and hdsEEG electrodes (6.5 ± 1.4 mm) were close in value, whereas the mean FWHM for theta for hdsEEG electrodes (17.3 ± 6.7 mm) was greater than FWHM for SDEs (13.0 ± 2.3 mm) and gray matter sEEGs (12.1 ± 1.8 mm).

### sEEG electrode location in white or gray matter influences FWHM measures

Within sEEG electrode pairs, signal redundancy between electrodes in gray matter was significantly decreased relative to their white matter located counterparts (Supplementary Figure 2). Signal attenuation is dependent on the conductivity ratio of the medium (Rogers et al. 2020), and white matter is considered largely anisotropic (Nunez and Srinivasan 2005), especially at this scale of field potential recording (Howell and McIntyre 2016). As such, white matter has been found to reflect activity from distant gray matter signals as well as volume conduction from nearby gray matter, thus increasing the likelihood of spurious correlation with activity in adjacent or distant regions (Mercier et al. 2017). While average FWHM was significantly greater for white matter located sEEGs (BHG: 11.3 ± 3.2 mm) than gray matter located sEEGs (BHG: 7.14 ± 1.7 mm), the average power of activity recorded at white matter located sEEG electrodes was significantly lower than gray matter located electrodes (Supplementary Figure 3). This is consistent with previous findings that electrodes located farther from gray matter signal generators record lower amplitude signals (Mercier et al. 2017; Young et al. 2019). As such, the current analyses comparing FWHM across electrode type, referencing scheme, and frequency spectra considered only gray matter located sEEG electrodes to avoid confounds in measures of correlation over distance due to signal attenuation.

### Impact of referencing schema on FWHM measures

Referencing schemes have an often-understated impact on signal detection, and how the data are referenced is a critical consideration in analyses of neural data (Li et al. 2018). The process of referencing neural signals has been found to distort and artificially inflate neural activation, functional connectivity and other measures (Li et al. 2018; Liu et al. 2015; Mercier et al. 2017). While measures of correlation should be scale-independent, the process of re-referencing likely influences correlation measures due to a decrease in distant noise, aiding in improved signal to noise ratio between nearby electrode pairs (Hnazaee et al. 2020). In our data, referencing scheme did not significantly influence FWHM measures for SDE or hdsEEG electrode pairs. However, for sEEG electrode pairs, we found a significant effect of referencing scheme on FWHM measures (Figure 4D,E). We found the choice of bipolar referencing scheme generates significantly lower FWHM measures between proximal sEEG electrode pairs, as compared with CAR and white matter referencing. These results corroborate previous findings (Li et al. 2018) comparing the effect of referencing method on Pearson’s correlation values averaged across sEEG electrode pairs regardless of inter-electrode distance.

While common average referencing is commonly implemented in icEEG analyses, there are many considerations when implementing a bipolar referencing scheme (Li et al. 2018; Mercier et al. 2017). While bipolar referencing removes all signal common to neighboring electrodes, this does not take into account anatomical location or dipole orientation, which can distort source localization (Hu et al. 2010). Depending on the location and orientation of sEEG electrodes relative to sulci and sources, bipolar referencing could have quite a variable effect on signal detection. Additionally, it is common when analyzing icEEG datasets to combine activity recorded via SDE and sEEG electrodes. In this case, the question of how to implement bipolar referencing in SDE electrodes becomes geometrically complex. As such, the FWHM for sEEG electrode pairs under the CAR scheme is found to reflect a very local listening zone (7.14 ± 1.7 mm), and these data suggest referencing scheme is a critical consideration in ensuring common noise to all electrodes is eliminated and does not confound further analysis.

### Comparison with previous studies

From neuroscientific research to the continuing development of brain-computer interfaces, decoding neural activity remains a necessary and complex goal. As neural interfaces continue to develop and our ability to record electrical activity from the brain at smaller scales advances, the overlap between what is feasible and what is informative remains unclear. A pivotal question of optimizing coverage, recording scale, or inter-electrode distance when designing neural interfaces remains a critical constraint. Thus, optimal balance between electrode type, size, and spacing of contacts will improve comprehensive mapping of cortical activity while minimizing redundancy of information.

Determining the optimal spacing and location of electrodes to not only minimize signal redundancy, but to also capture separable field potential recordings represents a pivotal hurdle for invasive field potential recordings in humans (Cybulski et al. 2015). There is currently no consensus in how to best allocate activity recorded by various electrode types to regions of nearby cortical space. In the absence of a solution to this problem, various methods are used to estimate the spatial extent of the neural population contributing to activity recorded at an individual electrode. The current methodologies implemented rely on assumptions and in vivo measurements to model the dielectric, conductive, and anisotropic aspects of neural tissue (Howell and McIntyre 2016, 2017; Miceli et al. 2017). These include spatial discrimination techniques (Herreras 2016), surface-based estimates of the recording zone (Kadipasaoglu et al. 2015) and weighting functions based on electrode properties of size, layout, and impedance (Dubey and Ray 2019). Computational models incorporating heterogeneity and anisotropy have been found to more accurately reconstruct neural response to stimulation in DBS application (Åström et al. 2012; Howell and McIntyre 2017).

In non-human primates, concurrent comparison of field potential recordings with single-unit (Dubey and Ray 2019) and multi-unit (Xing et al. 2009) resolution reveals the estimated spatial spread of cortical field potential recordings using intracranial microelectrodes (1 mm long; 400 μm pitch) to be local (roughly 3 mm) (Dubey and Ray 2020). In contrast, the location and design of neural probes in humans are largely limited to clinical application, making confident parameterization difficult. Despite these limitations, previous research has compared recording scale in humans (Halgren et al. 2018; Kellis et al. 2016; Lai et al. 2018; Muller et al. 2016; Trumpis et al. 2021) in order to disambiguate the uncertain properties of neural activity captured by different electrodes.

Modern icEEG recordings incorporate data from varying recording scales, cortical locations, referencing strategies, and analysis approaches. There is a wealth of existing data that has been gathered with a variety of tools and methodologies; the question becomes, how can findings be integrated across this diversity of scales? Beyond human neuroscience, how can direct comparisons be made with data collected from non-human primates? While icEEG recordings provide unique and robust high spatial and temporal resolution neural data, there are such disparate values of the spatial extent of LFP values reported in the literature (Kajikawa and Schroeder 2011; Kellis et al. 2016).

### Conclusions

Our results implicate electrode spacing, location, referencing strategy, and spectral band to be pivotal considerations in the minimization of signal redundancy and other confounds influencing the clarity of field potential analyses. We explored these confounds in a large robust dataset to probe these intrinsic uncertainties of field potential recordings. As with all aspects of scientific research, it is only through understanding the limitations of the tools we have to observe neural phenomenon that we can optimize the strengths, and get closer to understanding complex aspects of human cognition.

## Acknowledgements

We express our gratitude to all the patients who participated in this study; the neurologists at the Texas Comprehensive Epilepsy Program who participated in the care of these patients; and the nurses and technicians in the Epilepsy Monitoring Unit at Memorial Hermann Hospital who helped make this research possible. This work was supported by the NIH U01 N5098981, NIH/NIDCD DC014589, and the University of Texas System funding for the Texas Institute for Restorative Neurotechnologies.

## Supplemental Figures

**Supplementary Figure 1.**
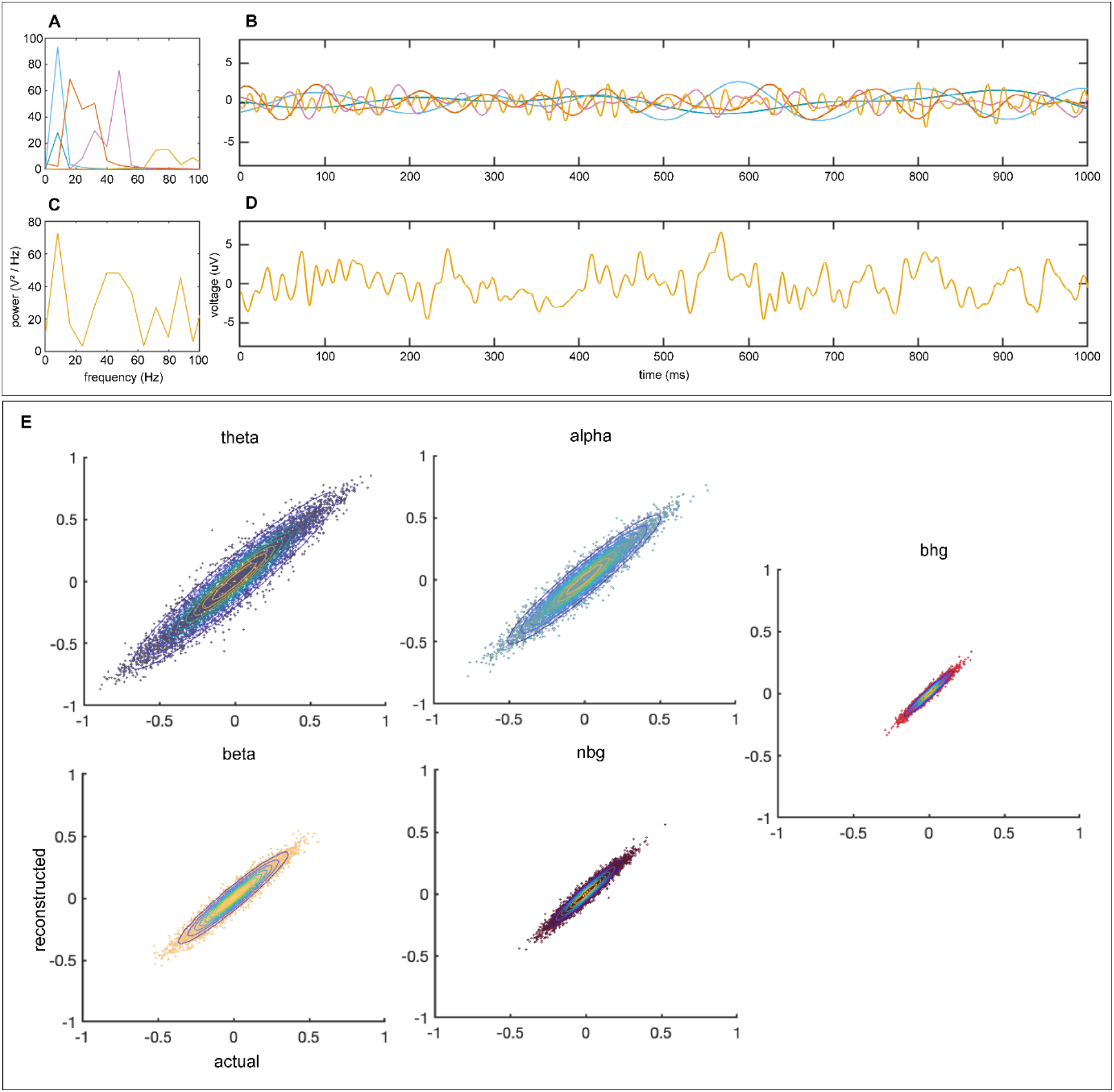
Correlation analysis on simulated timeseries data reveals no spurious correlation due to the analytic pipeline. Representative spectral power (**A**) and timeseries (**B**) of simulated neural data in each frequency range of interest (Theta, 4-8 Hz; Alpha, 8-15 Hz; Beta, 15-30 Hz; Narrowband Gamma, 30-60 Hz; Broadband High Gamma, 70-150 Hz). Representative power spectrum (**C**) and timeseries (**D**) of electric field signal comprised of summed timeseries in each frequency shown in (**B**). Comparison of actual and reconstructed Pearson’s correlation coefficient (r) between every combination of simulated timeseries (**E**) overlayed with 2D probability density estimation reveal no significant difference between actual and reconstructed correlation values on simulated data.

**Supplementary Figure 2.**
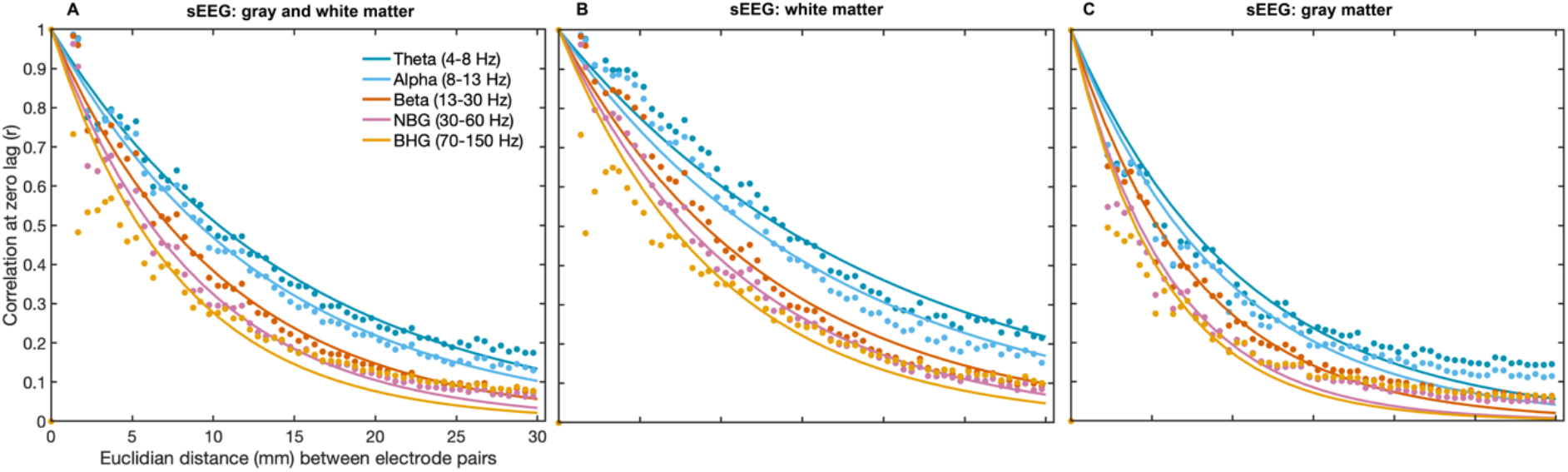
Anatomical location of sEEG contacts in gray or white matter significantly influences correlation measures over distance. Pearson’s correlation coefficient was measured between pairs of sEEG electrodes located in gray and white matter (**A**; 47 patients; 6,757 electrodes; 244,621 electrode pairs), white matter only (**B**; 2,649 electrodes; 43,957 electrode pairs), or gray matter only (**C**; 2,916 electrodes; 47,522 electrode pairs). Each data point is binned into 0.5 mm bins based on distance between electrode pairs, colored based on frequency range of interest and fit with an exponential decay function shown as colored solid lines. Abbreviations: narrowband gamma (NBG), broadband high gamma (BHG).

**Supplementary Figure 3.**
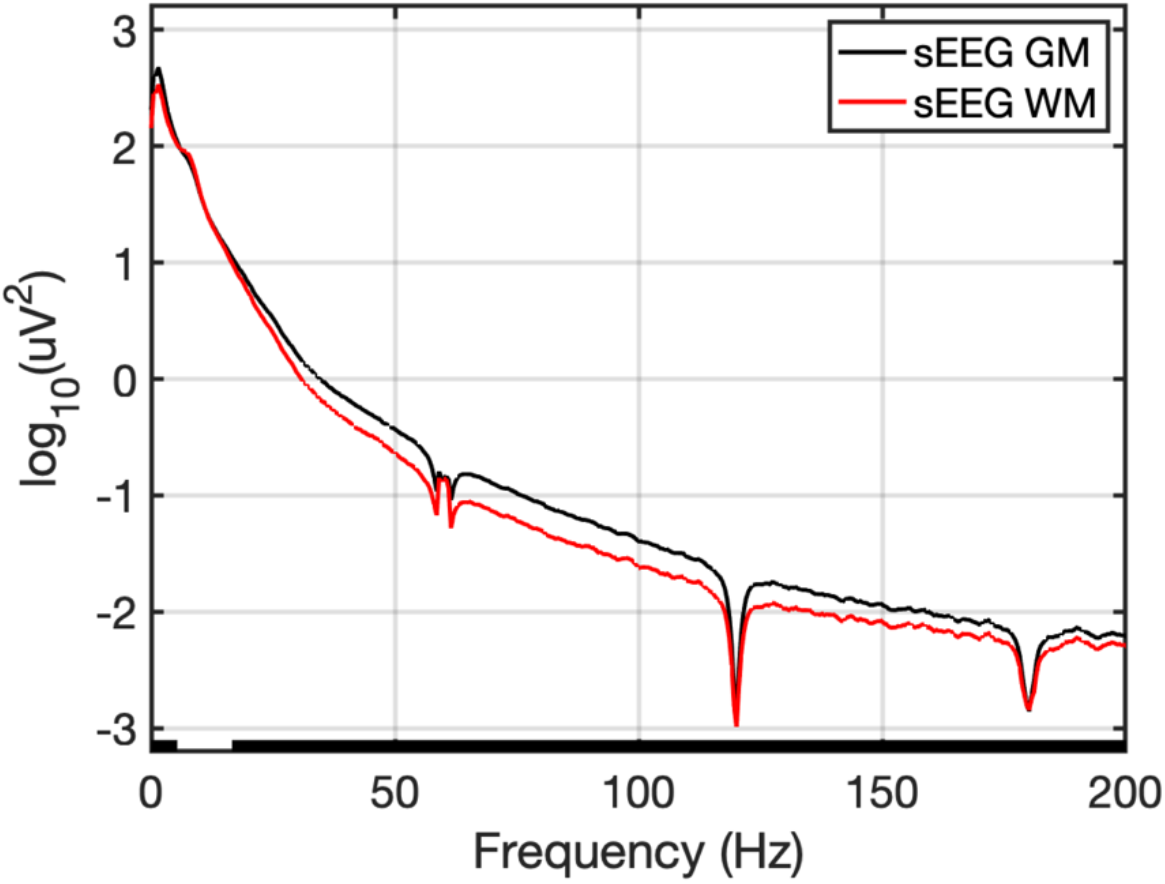
Mean power over frequency for sEEG electrodes based on location in gray or white matter. Mean power spectral density (PSD) plots for sEEG electrodes located in white matter (WM; red; 2,649 electrodes) or gray matter (GM; black; 2,916 electrodes). Notch filters were applied at 60 Hz and harmonics. Results from Wilcoxon sign rank test with significance threshold of <0.01 denoted by black bar along the x axis.

**Supplementary Figure 4.**
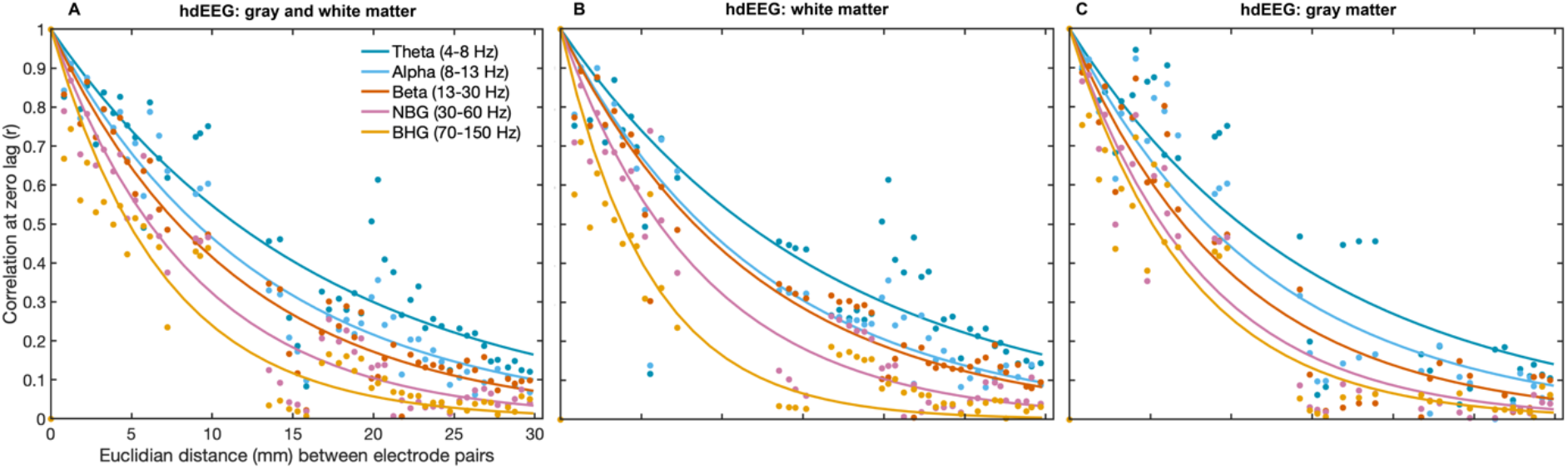
Impact of anatomical location of high-density sEEG (hdsEEG) electrodes on correlation measures over distance. Pearson’s correlation coefficient was measured between pairs of hdsEEG electrodes located in gray and white matter (**A**; 6 patients; 153 electrodes; 1,967 electrode pairs), white matter only (**B**; 53 electrodes; 421 electrode pairs), or gray matter only (**C**; 59 electrodes; 347 electrode pairs). Each data point is the correlation values for each patient binned into 0.5 mm bins based on distance between electrode pairs, colored based on frequency range of interest and fit with an exponential decay function shown as colored solid lines. Abbreviations: narrowband gamma (NBG), broadband high gamma (BHG).

## References

Allen M, Poggiali D, Whitaker K, Marshall TR, Kievit RA. Raincloud plots: a multi-platform tool for robust data visualization. Wellcome Open Res 4: 63, 2019.

Åström M, Lemaire J-J, Wårdell K. Influence of heterogeneous and anisotropic tissue conductivity on electric field distribution in deep brain stimulation. Med Biol Eng Comput 50: 23–32, 2012.

Bartoli E, Conner CR, Kadipasaoglu CM, Yellapantula S, Rollo MJ, Carter CS, Tandon N. Temporal Dynamics of Human Frontal and Cingulate Neural Activity During Conflict and Cognitive Control. Cereb Cortex New York N Y 1991 28: 3842–3856, 2018.

Bingham CS, Paknahad J, Girard CBC, Loizos K, Bouteiller J-MC, Song D, Lazzi G, Berger TW. Admittance Method for Estimating Local Field Potentials Generated in a Multi-Scale Neuron Model of the Hippocampus. Front Comput Neurosc 14: 72, 2020.

Buzsáki G, Anastassiou CA, Koch C. The origin of extracellular fields and currents — EEG, ECoG, LFP and spikes. Nat Rev Neurosci 13: 407–420, 2012.

Cogan GB, Thesen T, Carlson C, Doyle W, Devinsky O, Pesaran B. Sensory–motor transformations for speech occur bilaterally. Nature 507: 94–98, 2014.

Cole S, Donoghue T, Gao R, Voytek B. NeuroDSP: A package for neural digital signal processing. J Open Source Softw 4: 1272, 2019.

Conner CR, Ellmore TM, DiSano MA, Pieters TA, Potter AW, Tandon N. Anatomic and electro-physiologic connectivity of the language system: a combined DTI-CCEP study. Comput Biol Med 41: 1100–9, 2011.

Conner CR, Kadipasaoglu CM, Shouval HZ, Hickok G, Tandon N. Network dynamics of Broca’s area during word selection. Plos One 14: e0225756, 2019.

Cox RW. AFNI: Software for Analysis and Visualization of Functional Magnetic Resonance Neuroimages. Comput Biomed Res 29: 162–173, 1996.

Cybulski TR, Glaser JI, Marblestone AH, Zamft BM, Boyden ES, Church GM, Kording KP. Spatial information in large-scale neural recordings. Front Comput Neurosc 8: 172, 2015.

Dale AM, Fischl B, Sereno MI. Cortical Surface-Based Analysis I. Segmentation and Surface Reconstruction. Neuroimage 9: 179–194, 1999.

Derner M, Chaieb L, Surges R, Staresina BP, Fell J. Modulation of Item and Source Memory by Auditory Beat Stimulation: A Pilot Study With Intracranial EEG. Front Hum Neurosci 12: 500, 2018.

Dubey A, Ray S. Cortical Electrocorticogram (ECoG) Is a Local Signal. J Neurosci 39: 4299–4311, 2019.

Dubey A, Ray S. Comparison of tuning properties of gamma and high-gamma power in local field potential (LFP) versus electrocorticogram (ECoG) in visual cortex. Sci Rep-uk 10: 5422, 2020.

Ellenrieder N von, Khoo HM, Dubeau F, Gotman J. What do intracerebral electrodes measure? Clin Neurophysiol 132: 1105–1115, 2021.

Forseth KJ, Kadipasaoglu CM, Conner CR, Hickok G, Knight RT, Tandon N. A lexical semantic hub for heteromodal naming in middle fusiform gyrus. Brain J Neurology 141: 2112–2126, 2018.

Foster BL, Dastjerdi M, Parvizi J. Neural populations in human posteromedial cortex display opposing responses during memory and numerical processing. Proc National Acad Sci 109: 15514–15519, 2012.

Guillory SA, Bujarski KA. Exploring emotions using invasive methods: review of 60 years of human intracranial electrophysiology. Soc Cogn Affect Neur 9: 1880–1889, 2014.

Halgren M, Fabó D, Ulbert I, Madsen JR, Erőss L, Doyle WK, Devinsky O, Schomer D, Cash SS, Halgren E. Superficial Slow Rhythms Integrate Cortical Processing in Humans. Sci Rep-uk 8: 2055, 2018.

Herreras O. Local Field Potentials: Myths and Misunderstandings. Front Neural Circuit 10: 101, 2016.

Hnazaee MF, Wittevrongel B, Khachatryan E, Libert A, Carrette E, Dauwe I, Meurs A, Boon P, Roost DV, Hulle MMV. Localization of deep brain activity with scalp and subdural EEG. Neuroimage 117344, 2020.

Howell B, McIntyre CC. Analyzing the tradeoff between electrical complexity and accuracy in patient-specific computational models of deep brain stimulation. J Neural Eng 13: 036023, 2016.

Howell B, McIntyre CC. Role of Soft-Tissue Heterogeneity in Computational Models of Deep Brain Stimulation. Brain Stimul 10: 46–50, 2017.

Hu S, Stead M, Dai Q, Worrell GA. On the Recording Reference Contribution to EEG Correlation, Phase Synchorony, and Coherence. Ieee Transactions Syst Man Cybern Part B Cybern 40: 1294–1304, 2010.

Kadipasaoglu CM, Forseth K, Whaley M, Conner CR, Rollo MJ, Baboyan VG, Tandon N. Development of grouped icEEG for the study of cognitive processing. Front Psychol 6: 1008, 2015.

Kajikawa Y, Schroeder CE. How Local Is the Local Field Potential? Neuron 72: 847–858, 2011.

Kellis S, Sorensen L, Darvas F, Sayres C, O’Neill K, Brown RB, House P, Ojemann J, Greger B. Multi-scale analysis of neural activity in humans: Implications for micro-scale electrocorticography. Clin Neurophysiol 127: 591–601, 2016.

Lachaux J-P, Axmacher N, Mormann F, Halgren E, Crone NE. High-frequency neural activity and human cognition: Past, present and possible future of intracranial EEG research. Prog Neurobiol 98: 279–301, 2012.

Lai M, Demuru M, Hillebrand A, Fraschini M. A comparison between scalp- and source-reconstructed EEG networks. Sci Rep-uk 8: 12269, 2018.

Łęski S, Lindén H, Tetzlaff T, Pettersen KH, Einevoll GT. Frequency Dependence of Signal Power and Spatial Reach of the Local Field Potential. Plos Comput Biol 9: e1003137, 2013.

Li G, Jiang S, Paraskevopoulou SE, Wang M, Xu Y, Wu Z, Chen L, Zhang D, Schalk G. Optimal referencing for stereo-electroencephalographic (SEEG) recordings. NeuroImage 183: 327–335, 2018.

Liu Y, Coon WG, Pesters A de, Brunner P, Schalk G. The effects of spatial filtering and artifacts on electrocorticographic signals. J Neural Eng 12: 056008, 2015.

Marblestone AH, Zamft BM, Maguire YG, Shapiro MG, Cybulski TR, Glaser JI, Amodei D, Stranges PB, Kalhor R, Dalrymple DA, Seo D, Alon E, Maharbiz MM, Carmena JM, Rabaey JM, Boyden ES, Church GM, Kording KP. Physical principles for scalable neural recording. Front Comput Neurosc 7: 137, 2013.

Martin AB, Yang X, Saalmann YB, Wang L, Shestyuk A, Lin JJ, Parvizi J, Knight RT, Kastner S. Temporal Dynamics and Response Modulation across the Human Visual System in a Spatial Attention Task: An ECoG Study. J Neurosci 39: 333–352, 2019.

Mercier MR, Bickel S, Megevand P, Groppe DM, Schroeder CE, Mehta AD, Lado FA. Evaluation of cortical local field potential diffusion in stereotactic electro-encephalography recordings: A glimpse on white matter signal. NeuroImage 147: 219–232, 2017.

Miceli S, Ness TV, Einevoll GT, Schubert D. Impedance Spectrum in Cortical Tissue: Implications for Propagation of LFP Signals on the Microscopic Level. Eneuro 4: ENEURO.0291-16.2016, 2017.

Miller CA, Behroozmand R, Etler CP, Nourski KV, Reale RA, Oya H, Kawasaki H, Greenlee JDW. Neural correlates of vocal auditory feedback processing: Unique insights from electrocorticography recordings in a human cochlear implant user. Eneuro 8: ENEURO.0181-20.2020, 2021.

Muller L, Hamilton LS, Edwards E, Bouchard KE, Chang EF. Spatial resolution dependence on spectral frequency in human speech cortex electrocorticography. J Neural Eng 13: 056013, 2016.

Nunez PL, Srinivasan R. Electric Fields of the Brain. The Neurophysics of EEG. second. Oxford University Press, 2005.

Parvizi J, Kastner S. Promises and limitations of human intracranial electroencephalography. Nat Neurosci 21: 474–483, 2018.

Pasley BN, David SV, Mesgarani N, Flinker A, Shamma SA, Crone NE, Knight RT, Chang EF. Reconstructing Speech from Human Auditory Cortex. Plos Biol 10: e1001251, 2012.

Pesaran B, Vinck M, Einevoll GT, Sirota A, Fries P, Siegel M, Truccolo W, Schroeder CE, Srinivasan R. Investigating large-scale brain dynamics using field potential recordings: analysis and interpretation. Nat Neurosci 21: 903–919, 2018.

Pieters TA, Conner CR, Tandon N. Recursive grid partitioning on a cortical surface model: an optimized technique for the localization of implanted subdural electrodes: Clinical article. J Neurosurg 118: 1086–1097, 2013.

Rogers N, Thunemann M, Devor A, Gilja V. Impact of Brain Surface Boundary Conditions on Electrophysiology and Implications for Electrocorticography. Front Neurosci-switz 14: 763, 2020.

Rollo PS, Rollo MJ, Zhu P, Woolnough O, Tandon N. Oblique trajectory angles in robotic stereo-electroencephalography. J Neurosurg 1–10, 2020.

Rouse AG, Williams JJ, Wheeler JJ, Moran DW. Spatial co-adaptation of cortical control columns in a micro-ECoG brain–computer interface. J Neural Eng 13: 056018, 2016.

Salari E, Freudenburg ZV, Branco MP, Aarnoutse EJ, Vansteensel MJ, Ramsey NF. Classification of Articulator Movements and Movement Direction from Sensorimotor Cortex Activity. Sci Rep-uk 9: 14165, 2019.

Tandon N. Mapping of human language. In: Clinical Brain Maping, edited by Yoshor D, Mizrahi E. McGraw Hill Education, 2012, p. 203–218.

Tandon N, Tong BA, Friedman ER, Johnson JA, Allmen GV, Thomas MS, Hope OA, Kalamangalam GP, Slater JD, Thompson SA. Analysis of Morbidity and Outcomes Associated With Use of Subdural Grids vs Stereoelectroencephalography in Patients With Intractable Epilepsy. Jama Neurol 76: 672–681, 2019.

Tong BA, Esquenazi Y, Johnson J, Zhu P, Tandon N. The Brain is Not Flat: Conformal Electrode Arrays Diminish Complications of Subdural Electrode Implantation, A Series of 117 Cases. World Neurosurg 144: e734–e742, 2020.

Torres D, Makarova J, Ortuño T, Benito N, Makarov VA, Herreras O. Local and Volume-Conducted Contributions to Cortical Field Potentials. Cereb Cortex 29: 5234–5254, 2019.

Trumpis M, Chiang C-H, Orsborn AL, Bent B, Li J, Rogers JA, Pesaran B, Cogan G, Viventi J. Sufficient sampling for kriging prediction of cortical potential in rat, monkey, and human ECoG. J Neural Eng 18: 036011, 2021.

Xing D, Yeh C-I, Shapley RM. Spatial Spread of the Local Field Potential and its Laminar Variation in Visual Cortex. J Neurosci 29: 11540–11549, 2009.

Young JJ, Friedman JS, Panov F, Camara D, Yoo JY, Fields MC, Marcuse LV, Jette N, Ghatan S. Quantitative Signal Characteristics of Electrocorticography and Stereoelectroencephalography: The Effect of Contact Depth. J Clin Neurophysiol 36: 195–203, 2019.

